# Draft Genome Assembly of the Ancient Tetraploid Orphan Legume Marama Bean (*Tylosema esculentum*) with PacBio HiFi data

**DOI:** 10.1101/2023.03.16.532621

**Authors:** Jin Li, Christopher Cullis

## Abstract

*Tylosema esculentum* (marama bean), an underutilized orphan legume, has long been considered to have the potential to be domesticated as a crop to improve local food security due to the nutrient-rich seeds. As a plant species that grows naturally in the deserts of southern Africa, marama also serves as a good model for studying plant adaptation to extreme environments. In this study, HMW leaf DNA samples were prepared to generate 21.6 Gb PacBio HiFi data, which was assembled into to a raw tetraploid genome assembly of 1.24 Gb using Canu and into a partially phased assembly of 564.8 Mb by Hifiasm. The N50 values were 1.28 Mb and 2.75 Mb, respectively, and the BUSCO completeness were all above 99%. Repeats were found to account for 27.35% of the genome. The k-mer analysis indicated that marama was likely to be an autotetraploid plant with an estimated haplotype genome size of only 277 Mb. The current assembly was aligned with the genome of *Bauhinia variegata*, the closest species to marama whose genome has been sequenced, with an overall alignment rate of only 20.36% indicating a significant divergence between the two. This is the first high-quality genome assembly of marama bean, albeit unphased and still fragmented. However, some of the long contigs, which can be close to half the chromosome length, can serve as good references for studying the genes underlying the traits of interest. This will greatly facilitate the molecular breeding of the bean.

## Introduction

*Tylosema esculentum* (marama bean), a long-lived perennial legume from southern Africa, has long been considered to have the potential to be domesticated as a crop (Jackson et al., 2009) (Figure 1). Marama has a unique drought avoidance strategy by growing tubers weighing up to 500 pounds to store water, enabling it to survive the prolonged hot and dry conditions of the Kalahari Desert (Figure 1D) (Keith and Renew, 1975; Cullis et al., 2018). The domestication of marama is thought to be able to improve local food security due to the high nutritional value of the edible seeds, whose protein and lipid contents are comparable to those of the commercial crops soybean and peanuts, respectively (Dakora, 2013; Omotayo and Aremu, 2021). A major obstacle to marama breeding is that it does not flower until at least the second year after planting, making traditional breeding inefficient. Studying the genetic diversity that exists in nature and utilizing molecular marker-assisted breeding strategies are considered good alternatives (Cullis et al., 2019; Hasan et al., 2019). The main goals of marama breeding include developing plants with an erect habit, which would facilitate seed harvesting in the field, and overcoming self-incompatibility, which would allow the development of inbred lines to accelerate the production of new varieties with favorable allelic combinations (Enciso-Rodríguez et al., 2019; Cullis et al., 2022). Having a high-quality genome assembly will undoubtedly provide a reference for these studies.

**Figure 1.**
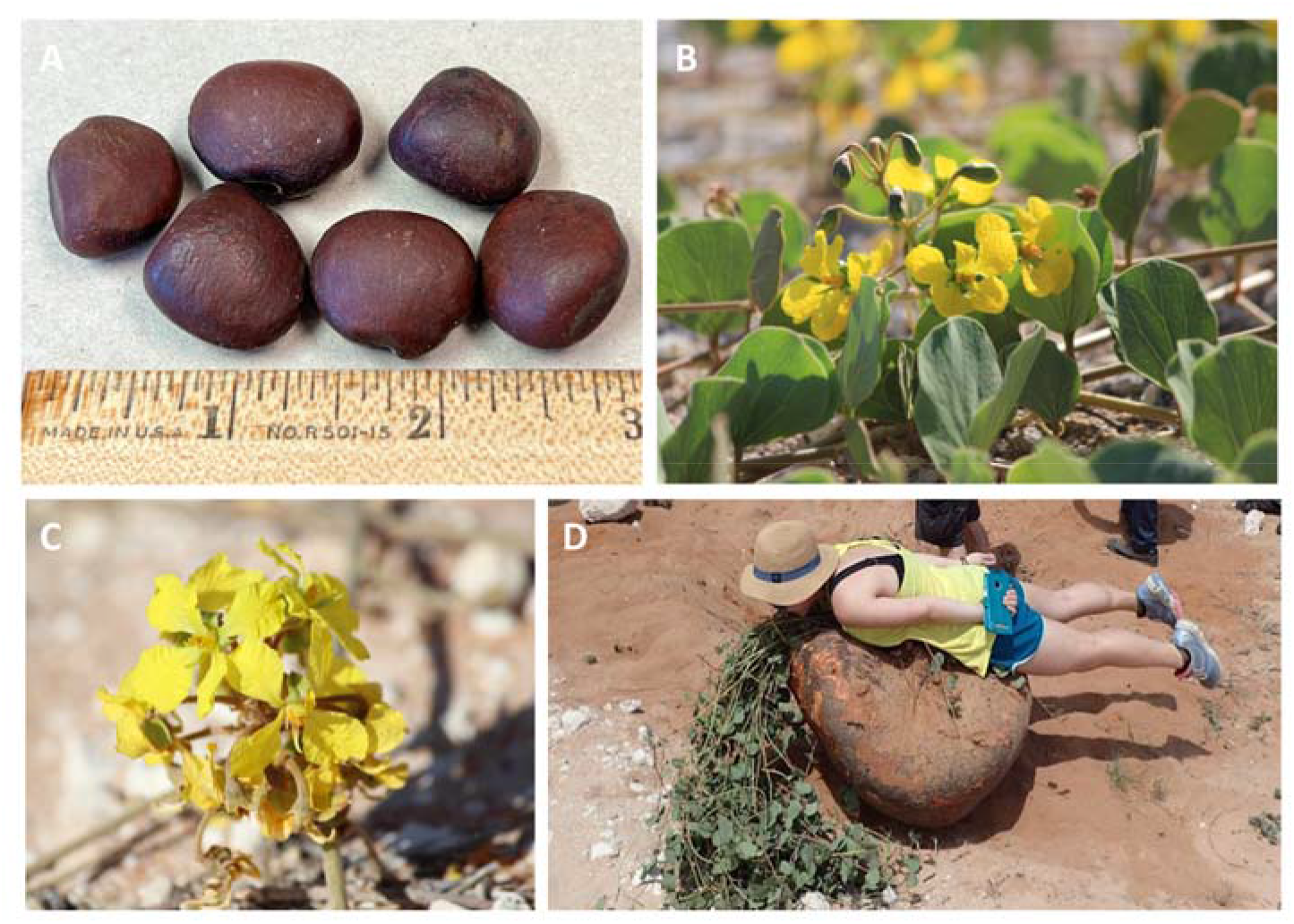
Morphology of wild *T. esculentum* (marama bean) from Namibia. (A). Brownish-black seeds, up to an inch long, are edible when roasted. The protein content is 30-39% dm and the lipid content is 35-48% dm (Amarteifio, 1998; Belitz et al., 2004). (B). Prostrate form with stems up to 3 m long (Bower et al., 1988) (C). Yellow flowers, beginning to bloom in midsummer for at least the second year after planting. (D). Giant tubers weighing over 500 pounds, 90% of the weight comes from water (Cullis et al., 2018).

The total genome size of *T. esculentum* was estimated to be 1 Gb with 44 chromosomes (2n=4x=44), according to the next-generation sequencing data and Feulgen staining (Takundwa et al., 2012; Cullis et al., 2019). Currently, Illumina whole genome sequencing data of more than 80 marama individuals collected from various geographical locations in Namibia and South Africa, as well as PacBio long reads from a few individuals are available and deposited under PRJNA779273. These have been successfully used in the assembly of marama chloroplast and mitochondrial reference genomes (Kim and Cullis, 2017; Li and Cullis, 2021). Comparative genomic analyses were also performed to investigate the genetic diversity present in the marama organelle genome (Li and Cullis, 2023). However, the assembly of the nuclear genome is still at a very rudimentary level, with an N50 value of only 3 kb, by Dr. Kyle Logue using only the Illumina reads of marama.

Genome assembly has been greatly facilitated as next-generation sequencing has become cheaper, faster, and with higher throughput (Von Bubnoff, 2008). However, for the assembly of complex genomes, including polyploid genomes and repeat-rich genomes, short reads generated by the next-generation sequencing cannot fulfill these tasks. As a third-generation sequencing technology, PacBio provides longer reads, with an average length over 10 kb and up to 25 kb, making up for the previous shortcomings. The latest PacBio HiFi sequencing improves the accuracy rate to over 99.9% on the basis of retaining the length of reads (Hon et al., 2020). In this study, the first high-quality genome assembly of marama was accomplished using only data from the PacBio HiFi platform assembled by the tools HiCanu and Hifiasm. This has particular significance in the projected molecular breeding research work on marama.

## Materials and Methods

### Sample collection and DNA extraction

Marama Sample 4 was an individual in the greenhouse of Case Western Reserve University grown from seeds collected in Namibia at undocumented location. 1 g of fresh young leaves were collected and ground thoroughly with a pestle in a mortar containing liquid nitrogen. DNA was then extracted using a Quick-DNA HMW MagBead kit (Zymo Research) following the protocol. Double-stranded DNA was quantified by the Invitrogen™ Qubit™ 3.0 Fluorometer after mixing 5 μl DNA with 195 μl working solution, and a 200 ng DNA sample was electrophoresed on a 1.5% agarose TBE gel at 40 V for 24 hours.

### Library preparation and sequencing

The prepared sample was sent to the Genomics Core Facility at the Icahn School of Medicine at Mount Sinai for PacBio sequencing. The HiFi sequencing library was prepared using the SMRTbell^®^ express template prep kit 2.0 (Pacific Biosciences, Menlo Park, CA, USA). SMRT sequencing was performed on two 8M SMRT^®^ Cells on the Sequel^®^ II system. 2,184,632 PacBio HiFi reads were generated with a total length of 21.6 Gb in the end. In addition, the Illumina WGS data of another 84 individuals (including samples M1, M40, Index1) were prepared previously and deposited in the NCBI SRA database (PRJNA779273) as described in the study (Li and Cullis, 2023).

### De novo genome assembly and evaluation

The 20.11 GB PacBio HiFi reads generated for *T. esculentum* Sample 4 were assembled using HiCanu (Nurk et al., 2020; https://canu.readthedocs.io/en/latest/quick-start.html#assembling-pacbio-hifi-with-hicanu) with input genome size set to 1 Gb according to the previous estimate (Cullis et al., 2019). Jellyfish 2.3.0 (Marçais and Kingsford, 2011; https://github.com/gmarcais/Jellyfish) was used to count k-mer for the PacBio HiFi reads of Sample 4 with the k-mer length set to 21 and to generate a k-mer count histogram, which was then used to draw k-mer spectra on GenomeScope 2.0 (Ranallo-Benavidez et al., 2020; http://qb.cshl.edu/genomescope/genomescope2.0/). The assembly quality was evaluated by QUAST 5.2.0. (Mikheenko et al, 2018; https://github.com/ablab/quast) and the results were visualized by Matplotlib v.1.3.1. (Hunter, 2007). The genome completeness was assessed by comparison with the Embryophyta ortholog database (embryophyta_odb10) containing 1614 genes and the Fabales ortholog database (fabales_odb10) containing 5366 genes using BUSCO v5.4.4 (Simão et al., 2015; https://busco.ezlab.org/). Genome repeat annotation was conducted by RepeatMasker 4.1.4 (Tarailo-Graovac and Chen, 2009; https://www.repeatmasker.org/) with the Repbase library (Bao et al., 2015). The marama genome assembly was mapped to the haplotype genome assembly of *Bauhinia variegata* ASM2237911v2 (Zhong et al., 2022) via minimap2 v2.24 (Li, 2018; https://github.com/lh3/minimap2). The resulting pairwise mapping format (PAF) data was visualized by a dot plot drawn by the R package pafr (https://github.com/dwinter/pafr).

## Results

21.58 Gbp PacBio HiFi reads were obtained, based on which GenomeScope 2.0 was used to construct the k-mer distribution map, showing three peaks at 1-fold, 2-fold, and 4-fold coverage respectively (Figure 2). The PacBio data was found to fit the tetraploid model best despite marama was initially thought to be an ancient hexaploid plant. In addition, the k-mer spectra showed high heterozygosity, with 2.2% of the genome being heterozygous. The frequency of aaab (1.410%) was found to be greater than that of aabb (0.498%), indicating that the individual studied was possibly an autotetraploid (Ranallo-Benavidez et al., 2020). The estimated genome size of *T. esculentum* was only 277.4 Mb, suggesting that marama has a compact genome, as reported for the legume *Amphicarpaea edgeworthii*. It has a genome size of 298.1 Mb and also has 11 chromosomes (2*n* = 22) (Liang et al., 2009; Liu et al., 2021).

**Figure 2.**
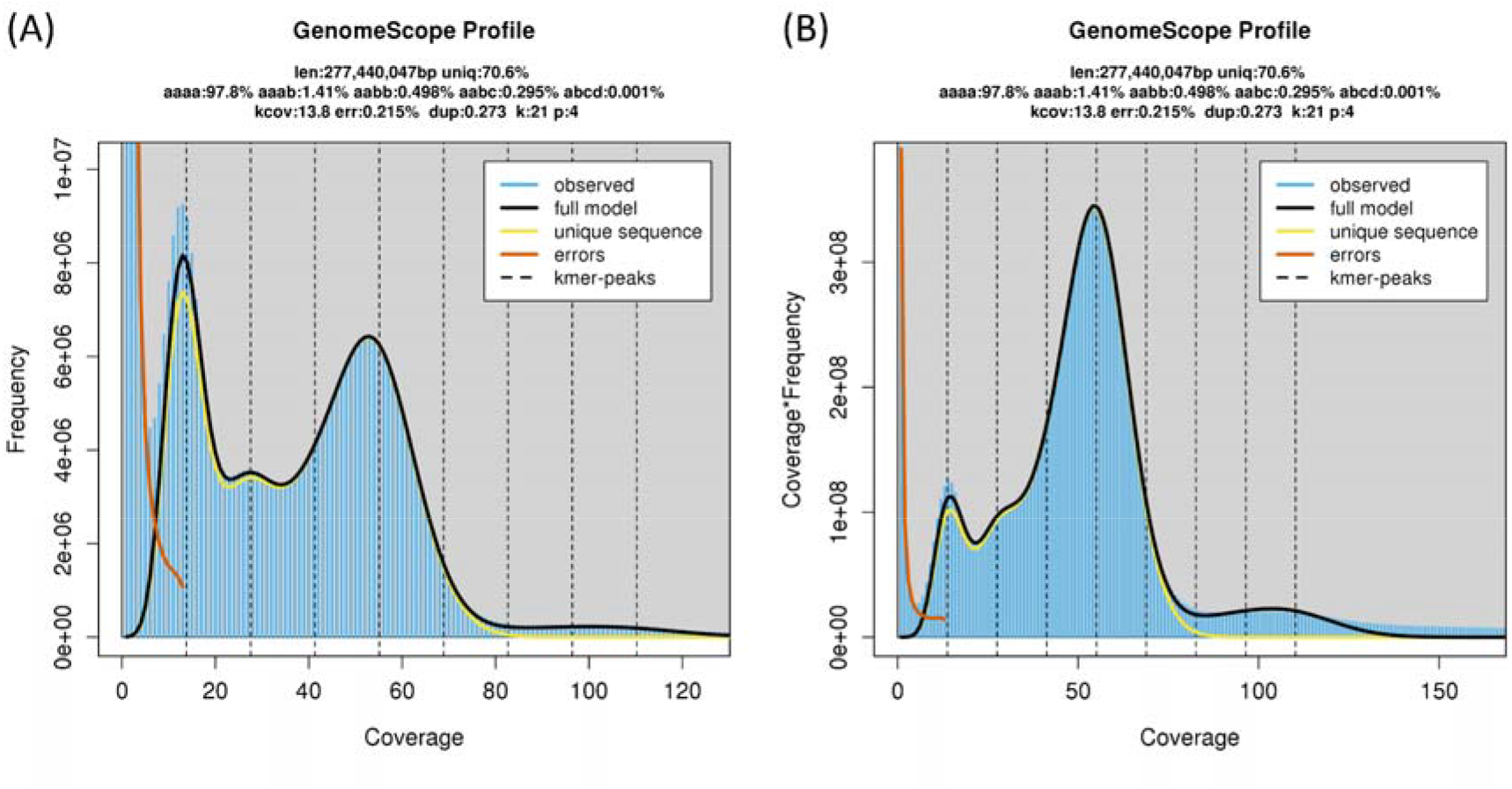
K-mer spectra built on the PacBio HiFi reads of Sample 4 using GenomeScope 2.0. (A). Frequency-coverage k-mer spectrum. (B). Coverage*frequency-coverage k-mer spectrum.

The 2,184,632 PacBio HiFi reads were assembled by Canu to generate a raw tetraploid genome assembly of 1.24 Gb consisting of 9532 contigs, with an N50 value of 1.28 Mb and an L50 of 252 (minimum number of contigs with a total length equal to half the genome size) (Table 1). The current genome assembly is considered to contain contigs from the four haplotype genomes in preparation for future haplotype phasing. The average length of the obtained contigs was 1.24 Mb, and the longest contig was 9.80 Mb (Figure 3A). The L90 value of the genome assembly was 3,235, which means that the total length of the top 3,235 contigs accounted for 90% of the genome size (Figure 3B). The average guanine-cytosine (GC) of all contigs was 36.06% (Table 1 and Figure 3C). BUSCO was used to evaluate the completeness of the *T. esculentum* genome assembly based on the comparison with 5366 genes from the Fabales ortholog database. 94.1% of these genes were found in the *T. esculentum* genome assembly. Furthermore, 99.5% of the genes from the Embryophyta ortholog database were detected in our assembly, indicating that it is highly complete. As expected, the proportions of duplicated genes were 93.6% and 98.7%, respectively, since most genes should have four copies in tetraploid plants. 27.35% of the *T. esculentum* genome assembly was annotated as repeats by RepeatMasker (Table 2). Long-terminal repeat (LTR) retroelements accounted for 13.21% of the genome, of which Ty1/Copia made up to 3.55% of the genome, and Gypsy/DIRS1 accounted for 9.14% of the genome. Low-complexity regions (LCRs) and simple sequence repeats (SSRs) were found to account for 6.47% and 4.05% of the *T. esculentum* genome size, respectively.

**Table 1.**
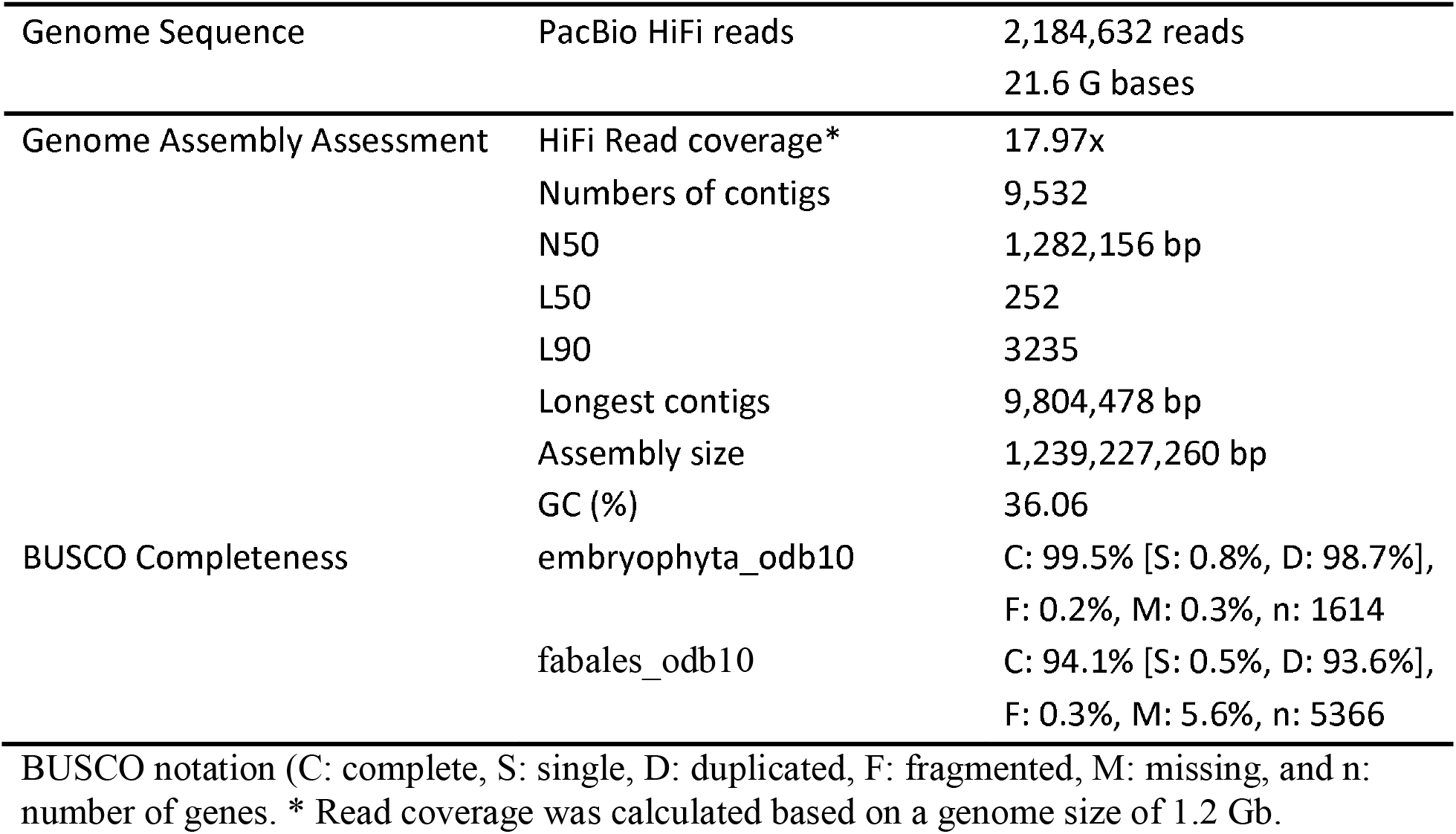
*T. esculentum* sequencing and draft genome assembly statistics

|  |  |  |
| --- | --- | --- |
| Genome Sequence | PacBio HiFi reads | 2,184,632 reads<br>21.6 G bases |
| Genome Assembly Assessment | HiFi Read coverage* | 17.97x |
|  | Numbers of contigs | 9,532 |
|  | N50 | 1,282,156 bp |
|  | L50 | 252 |
|  | L90 | 3235 |
|  | Longest contigs | 9,804,478 bp |
|  | Assembly size | 1,239,227,260 bp |
|  | GC (%) | 36.06 |
| BUSCO Completeness | embryophyta_odb10 | C: 99.5% [S: 0.8%, D: 98.7%],<br>F: 0.2%, M: 0.3%, n: 1614 |
|  | fabales_odb10 | C: 94.1% [S: 0.5%, D: 93.6%],<br>F: 0.3%, M: 5.6%, n: 5366 |

BUSCO notation (C: complete, S: single, D: duplicated, F: fragmented, M: missing, and n: number of genes. \* Read coverage was calculated based on a genome size of 1.2 Gb.

**Figure 3.**
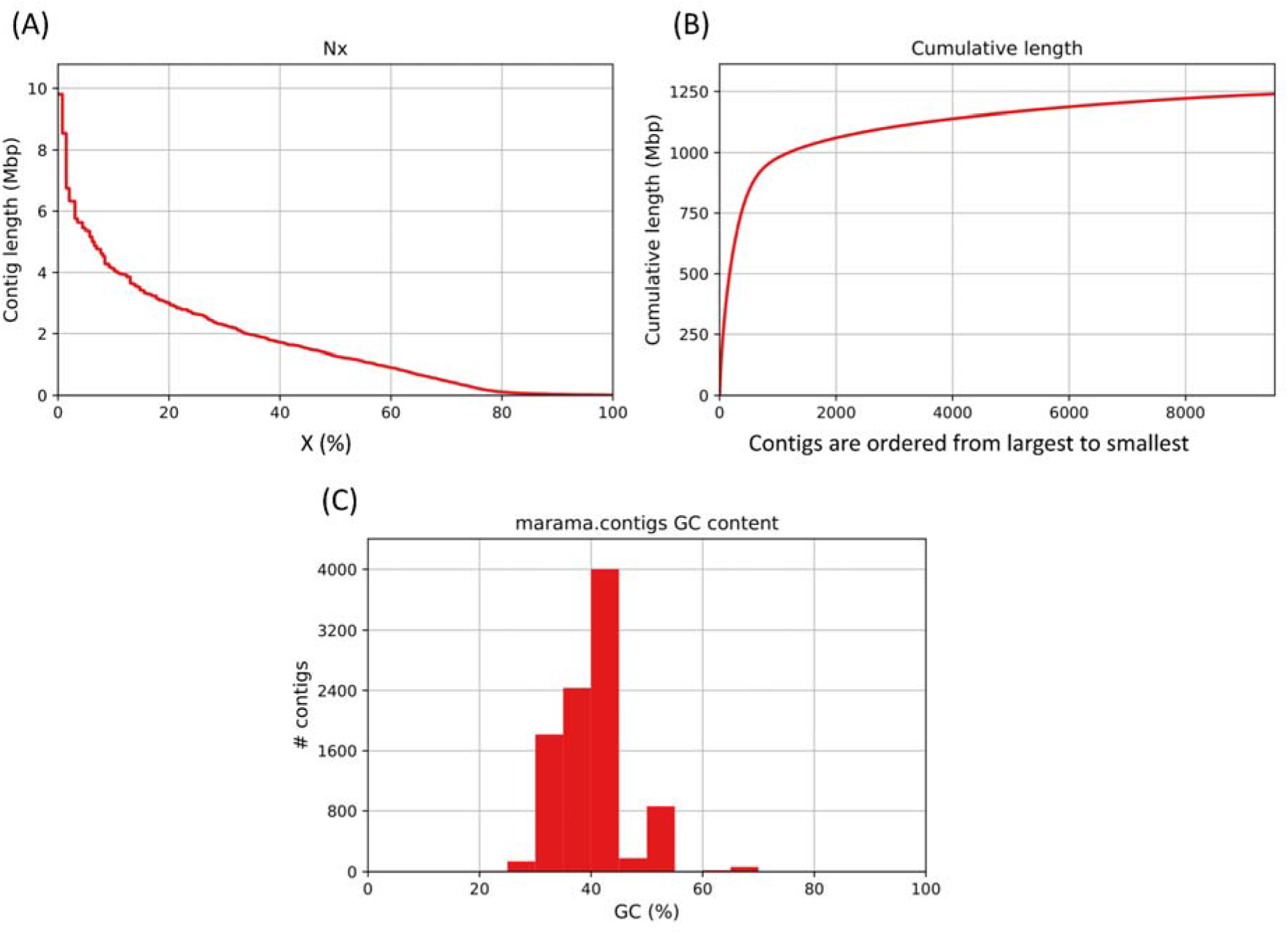
Genome assembly quality assessment plots drawn by QUAST 5.2.0. (A) Nx plot showing the distribution of contig lengths as x varies from 0 to 100%. (B) Cumulative length plot. The contigs were sorted from largest to smallest. (C) GC plot showing the distribution of GC content in the contigs.

**Table 2.** Summary of repeat elements in the *T. esculentum* genome assembly by RepeatMasker

|  | Number of elements* | Length occupied (bp) | Percentage of sequence (%) |
| --- | --- | --- | --- |
| Retroelements | 239,152 | 176,219,041 bp | 14.22% |
| SINEs: | 2,428 | 151,942 bp | 0.01% |
| Penelope | 4,697 | 257,720 bp | 0.02% |
| LINEs: | 56,298 | 12,306,565 bp | 0.99% |
| CRE/SLACS | 148 | 13,819 bp | 0.00% |
| L2/CR1/Rex | 3,208 | 171,686 bp | 0.01% |
| R1/LOA/Jockey | 1,800 | 96,061 bp | 0.01% |
| R2/R4/NeSL | 433 | 22,878 bp | 0.00% |
| RTE/Bov-B | 3,045 | 560,899 bp | 0.05% |
| L1/CIN4 | 39,173 | 10,918,951 bp | 0.88% |
| LTR elements | 180,426 | 163,760,534 bp | 13.21% |
| BEL/Pao | 2,400 | 286,196 bp | 0.02% |
| Ty1/Copia | 45,240 | 43,947,485 bp | 3.55% |
| Gypsy/DIRS1 | 111,682 | 113,301,860 bp | 9.14% |
| Retroviral | 5,724 | 287,611 bp | 0.02% |
| DNA transposons | 58,669 | 6,960,572 bp | 0.56% |
| hobo-Activator | 10,674 | 1,367,473 bp | 0.11% |
| Tc1-IS630-Pogo | 2,621 | 148,140 bp | 0.01% |
| En-Spm | 0 | 0 bp | 0.00% |
| MULE-MuDR | 10,709 | 1,298,115 bp | 0.10% |
| PiggyBac | 316 | 16,841 bp | 0.00% |
| Tourist/Harbinger | 3,996 | 537,089 bp | 0.04% |
| Other (Mirage, P-element, Transib) | 891 | 39,450 bp | 0.00% |
| Rolling-circles | 6,352 | 1,190,800 bp | 0.10% |
| Unclassified: | 3,289 | 278,950 bp | 0.02% |
| Total interspersed repeats: |  | 183,458,563 bp | 14.80% |
| Small RNA: | 14,055 | 14,349,721 bp | 1.16% |
| Satellites: | 19,870 | 9,567,228 bp | 0.77% |
| Simple repeats: | 421,704 | 50,213,770 bp | 4.05% |
| Low complexity: | 114,847 | 80,219,159 bp | 6.47% |

\* Most repeats fragmented by insertions or deletions have been counted as one element

Genomes of only a few plants from the Cercidoideae subfamily have been assembled, of which *B. variegata* is the evolutionarily closest to *T. esculentum* (Wunderlin, 2010). The haplotype genome assembly of *B. variegata* ASM2237911v2 has a size of 326.4 Mb, and contains 14 chromosomes (2*n* = 28) ranging in length from 18,256,449 bp to 27,622,603 bp (Zhong et al., 2022). The genome assembly of *T. esculentum* was mapped to the *B. variegata* genome by minimap2 and the result was visualized as a dot plot using R package pafr (Figure 4). Some contigs from the *T. esculentum* genome assembly reached half the chromosome length of *B. variegata* and exhibited a high degree of collinearity, confirming the reliability of our assembly. When Illumina reads from randomly selected samples (M1, M40, Index1) were mapped to the *B. variegata* genome via Bowtie2 v2.4.4 (Langmead and Salzberg, 2012; https://github.com/BenLangmead/bowtie2), the overall alignment rate was only around 20.36%, and the alignment rate with the genome of *Vigna radiata* PRJNA301363 was only 2.7% (Kang et al., 2014), indicating that *T. esculentum* has a highly divergent genome from these legumes.

**Figure 4.**
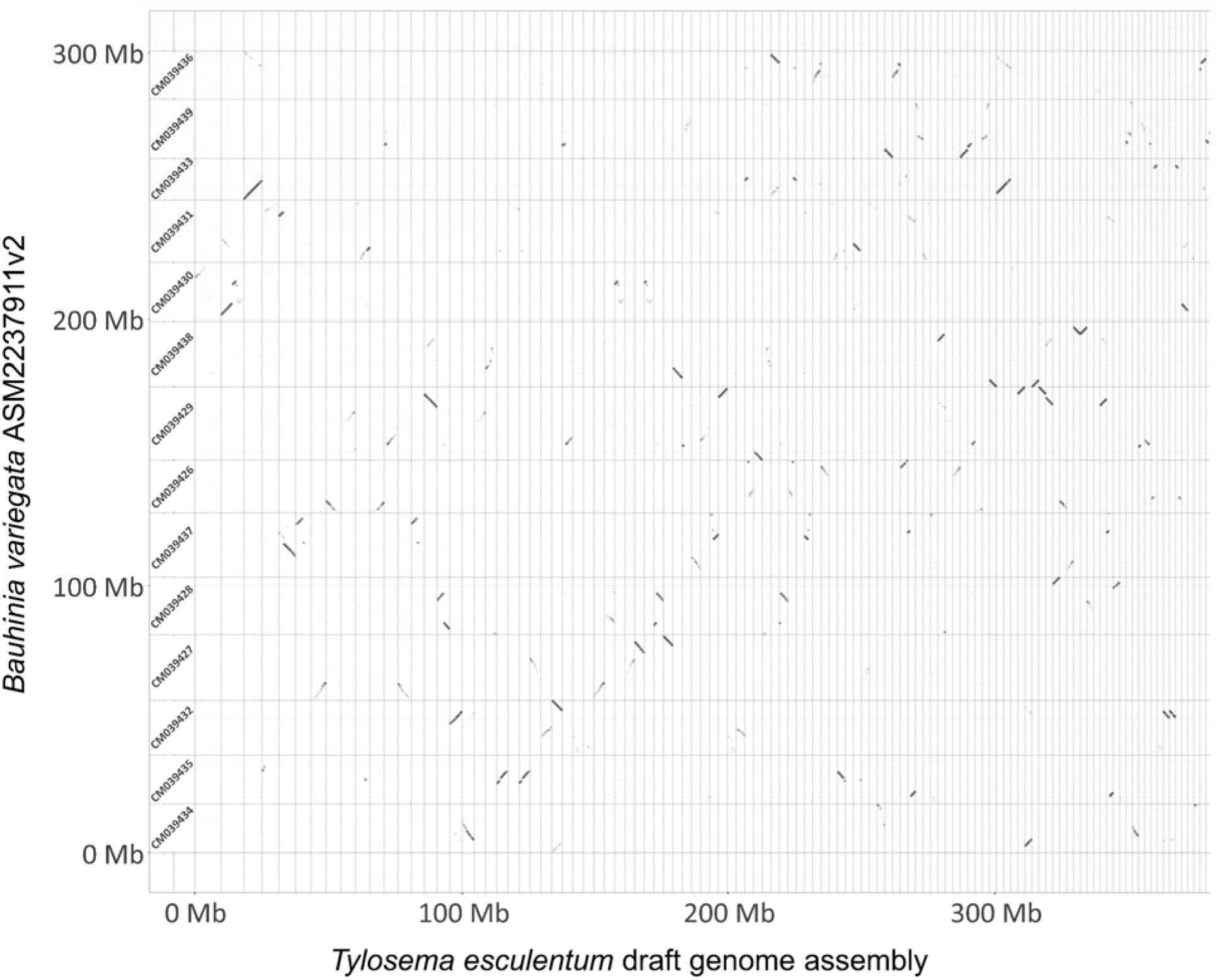
A dot plot of alignment of partial *T. esculentum* assembled contigs against the 14 chromosomes of *B. variegata* genome assembly ASM2237911v2. This figure was drawn by the R package pafr on the pairwise mapping format (PAF) document generated by minimap2. Each row represents one chromosome of the *B. variegata* genome with the chromosome GenBank ID labeled at the beginning of the row. Each column represents one contig from the *T. esculentum* genome assembly, sorted by size. Only the first 370 Mb of the 1.2 Gb assembly are included here, with highly fragmented contigs not shown. The black dotted lines show where the two genomes align. The ticks on both axes indicate the genomic scales in base pairs.

## Discussion

The is the first reported high-quality draft genome assembly of *T. esculentum* with an N50 value of 1.28 Mb, which has been dramatically improved from the 3 kb of the previous assembly done by Dr. Kyle Logue on Illumina reads. Although the current genome assembly of *T. esculentum* still contains numerous highly fragmented contigs, this is considered an ongoing project that will continual to be optimized in the future. Moreover, many of the obtained contigs are long enough to be used in the study of genes of interest, providing an important reference for marama breeding. The genomic resource lays a foundation for the study of genetic diversity existing in nature, which can be used to explore the genetic mechanism of self-incompatibility in marama and study the adaptation of plants to harsh environments, etc.

Another assembler Hifiasm 0.18.5 (Cheng et al., 2021; https://github.com/chhylp123/hifiasm) was used for haplotype assembly of *T. esculentum* and generated a partially phased assembly of 564.8 Mb. This contained 4,123 contigs with an N50 value of 2.75 Mb and an L50 of 35. The BUSCO score was 99.1% (S:62.0%, D:37.1%, F:0.4%, M:0.5%, n=1614) compared to the Embryophyta ortholog database (embryophyta_odb10) and 93.5% (S:45.5%, D:48.0%, F:0.5%, M:6.0%, n=5366) to the Fabales ortholog database (fabales_odb10). The contigs generated by Hifiasm have longer N50 and better continuity, but it is still a partially phased genome assembly that contains a large number of duplications, which need to be purged by third-party tools. However, this may also collapse repeats or segmental duplications that should have been included. Although the assembly from HiCanu is more fragmented, it is considered to have more complete genetic information because the default parameters of HiCanu separate haplotypes down to 0.01% divergence to generate a raw tetraploid assembly which avoid collapsing the genome. In the future, by using data from other sequencing platforms like Hi-C, contigs from the same chromosome can be grouped and then further scaffolded to the near-chromosome level (Aiden et al., 2009; Mascher et al., 2017).

## Acknowledgments

The authors would like to thank K. Logue for help with the initial genome assembly, to P. Chimwamurombe, M. Takundwa, J. Vorster, and K. Kunert for providing marama samples from Namibia and from the University of Pretoria Farm, to students in BIOL 301/401 in 2015 for assistance with DNA isolation and in BIOL 309 in 2018 for sample collection.

## Author Contributions

J.L. was involved in DNA extraction and carried out the genome assembly, data analysis and drafted the manuscript. C.C. conceived of the original idea for the project, provided extracted DNAs, supervised the research, and assisted in writing and editing the manuscript. All authors contributed to the article and approved the submitted version.

